# Out of sight, out of mind: Responses in primate ventral visual cortex track individual fixations during natural vision

**DOI:** 10.1101/2023.02.08.527666

**Authors:** Will Xiao, Saloni Sharma, Gabriel Kreiman, Margaret S. Livingstone

## Abstract

During natural vision, primates shift their gaze several times per second with large, ballistic eye movements known as saccades. Open questions remain as to whether visual neurons retain their classical retinotopic response properties during natural vision or whether neurons integrate information across fixations and predict the consequences of impending saccades. Answers are especially wanting for vision in complex scenes relevant to natural behavior. We let 13 monkeys freely view thousands of large natural images, recorded over 883 hours of neuronal responses throughout the ventral visual pathway across 4.7 million fixations, and designed flexible analyses to reveal the spatial, temporal, and feature selectivity of the responses. Ventral visual responses followed each fixation and did not become gaze-invariant as monkeys examined an image over seconds. Computational models revealed that neuronal responses corresponded to eye-centered receptive fields. The results suggest that ventral visual cortex remains predominantly retinotopic during natural vision and does not establish a gaze-independent representation of the world.

---

We see the world as stable, yet our eyes are in constant motion. How does the brain account for the motion of its visual sensor to construct a stable percept? The centuries-old question of visual stability dates back to von Helmholtz, Descartes, and Alhazen (Melcher, 2011; Wurtz, 2008). A proposed mechanism for visual stability is neurons that predictively remap the part of the visual field they represent. Studies have reported frontal, parietal, and superior collicular neurons that have retinotopic visual responses but respond to a stimulus displayed in their *future* receptive fields (RFs) before impending eye movements (Duhamel et al., 1992; Umeno & Goldberg, 1997; Walker et al., 1995). Such predictively remapping neurons are posited to contribute to visual stability by monitoring congruity between the pre- and post-saccadic scene (Wurtz, 2008).

Predictive remapping alone is insufficient to account for visual stability. Neurons that predictively shift their RFs only decouple two changes in time. It is unknown whether or which downstream neurons compare the predictive responses to postdictive ones to maintain a stable representation. Moreover, studies have probed remapped RFs with simple, often transient stimuli, such as a light spot, oriented bar, or grating, leaving it unknown whether remapped RFs exhibit any selectivity necessary for the stable representation of what is where. The processing of detailed visual form is specialized to the ventral visual pathway (Ungerleider, 1982), whose output could contain such a stable representation. Ventral visual processing culminates in the inferior temporal cortex (IT). IT is only two to three synapses away from the entorhinal cortex, which contains non-retinotopic gaze direction grid cells (Killian et al., 2012), and the hippocampus, which harbors place cells and consolidates episodic memory (Hartley et al., 2014; Rueckemann & Buffalo, 2017), plausibly with a gaze-invariant representation.

Most studies of ventral vision focus on the first-pass visual processing during stable fixation. Some previous investigations of ventral visual processing during eye movements have found neurons that predictively remap in V4 (Neupane et al., 2016; Tolias et al., 2001) and, to a lesser extent, in V1 and V2 (Nakamura & Colby, 2002). Others examined the feature selectivity of neurons in V1 and IT and found it to remain retinotopic (DiCarlo & Maunsell, 2000; Livingstone et al., 1996). DiCarlo and Maunsell further showed that IT responses had identical dynamics during free and passive viewing, providing evidence against predictive remapping in IT.

These studies continued to use simple stimuli in a sparse display. Intuitively, our experience of the world as stable may also rely on recurrent processing of a persistent scene rich in framing cues. Visual processing is affected by at least a few seconds of viewing history through adaptation (Solomon & Kohn, 2014). Active natural vision may recruit additional top-down modulation of visual processing (Gilbert & Li, 2013). A few studies have examined visual processing during free viewing of persisting, natural stimuli (McMahon et al., 2015; Podvalny et al., 2017; Rolls et al., 2003; Sheinberg & Logothetis, 2001) and have provided some evidence for non-classical responses. However, the interpretation of free-viewing data is hindered by challenges inherent in describing essentially single-trial neuronal responses in the face of unrestricted eye movements, complex stimuli, and intricate feature selectivity.

We studied monkeys freely viewing natural images and evaluated how traditional visual response properties manifested during this naturalistic behavior. We designed analysis methods flexible and robust enough to apply to a wide range of experimental conditions. Here, we report results summarized over 679 experimental sessions, containing 883 hours of recording from 13 monkeys making 4.7 million fixations on thousands of natural images. We found that responses along the ventral visual cortex primarily reflected the contents of the present fixation. The neuronal responses showed limited evidence of integrating the viewing history or predicting the consequence of upcoming saccades.

## Results

We recorded the activity of neurons throughout ventral visual cortex while monkeys freely viewed natural images. In each session, we presented a sequence of natural images that repeated in a pseudorandom block fashion (Fig. 1a). A trial lasted up to 1.5 s in a typical session (410 of 679 sessions; range 0.3–60 s in remaining sessions) and was interrupted if the monkey looked away from the image. Images were typically 16 × 16 degrees of visual angle (dva) in size (487 of 679 sessions; range 8 × 8–26 × 26 dva in remaining sessions). Monkeys naturally explored the images without explicit instructions or training. Across picture repeats, monkeys examined each picture with varied looking patterns (Fig. 1b inset). The basic statistics of the looking behavior were consistent across subjects (Fig. 1c–e) and matched previous studies on similar behaviors (Mitchell et al., 2014; Zhang et al., 2022). An average fixation lasted 248 ± 35 ms, while an average saccade took 48 ± 4 ms and subtended 4.9 ± 0.8 dva (all mean ± stdev. across subjects).

**Fig. 1.**
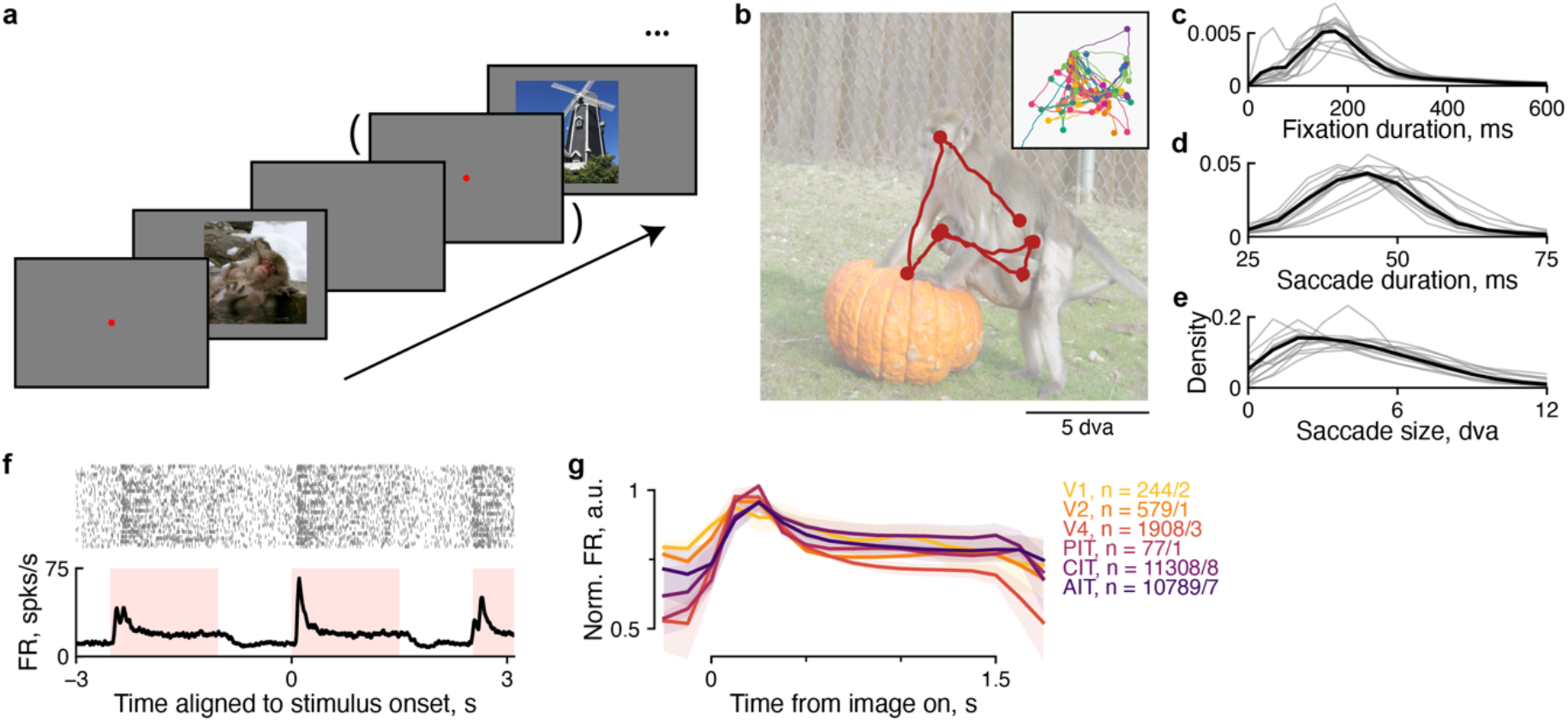
Overview of the free viewing experiment. **a**, The monkey freely viewed a random image in each trial. In a small subset of trials, a fixation dot was displayed before the image presentation. **b**, The eye trajectory in an example trial. The inset shows eye trajectories across trials for the same image in one experimental session. Colors indicate different trials; dots indicate fixations. **c–e**, Distribution of fixation durations, **c**, saccade durations, **d**, and saccade sizes, **e**. Each thin line indicates one monkey; the thick lines indicate the average across monkeys. **f**, Example spike rasters and average firing rates aligned to stimulus onset for an example neuron in anterior inferior temporal cortex (AIT). The pink shading indicates the stimulus presentation cadence in this session. **g**, Average normalized firing rates separately for each visual area. The n values indicate the total number of neurons/monkeys per visual area. Shading indicates the 95%-confidence interval (CI) of the mean by bootstrapping first over neurons, then over monkeys.

While monkeys freely viewed images, we recorded extracellular single- and multi-unit activity throughout the ventral visual pathway using chronically implanted multielectrode arrays. The recordings spanned six visual areas: V1, V2, V4, and IT in its posterior, central, and anterior subdivisions (PIT, CIT, and AIT). V1, V4, CIT, and AIT were represented by at least two monkeys. Overall, ventral visual neurons were more active during stimulus presentation than between trials (Fig. 1f–g). There was a transient increase in firing rate during stimulus onset, followed by sustained above-baseline firing while the monkeys looked around the image, returning to baseline after stimulus offset.

### Face-neuron responses were specific to each fixation

We started to examine how IT responses interacted with eye movements and stimulus content by focusing on recordings from face-selective neurons. Face neurons respond more strongly to faces than nonface objects presented during passive fixation (Tsao et al., 2006). Thus, we categorized fixations within 2.5 dva of a face region of interest (ROI) as face fixations and the rest as nonface fixations (Fig. 2a), assuming that IT neurons had roughly 5-dva receptive fields centered on the fovea. To identify face neurons functionally, we used the ‘zeroth fixation,’ the period right after image onset and before the first eye movement on the image. In this period, the initial presentation of a random image placed either a face or a nonface near where the monkey happened to be fixating. Thus, vision during the zeroth fixation was comparable to conventional passive viewing experiments. We calculated face selectivity indices (FSI; Methods) for neurons during zeroth fixations using the responses 70–200 ms after image onset. Face neurons were defined as units recorded from four face-patch arrays with FSI of at least 0.2, i.e., at least 50% higher responses to faces than nonfaces. Similarly, we calculated FSI during free viewing using responses 70–200 ms after the end of a saccade, i.e., the onset of non-zeroth fixations (‘fixation 1+’). Neurons were equally selective to faces during the zeroth and non-zeroth fixations (Fig. 2b).

**Fig. 2.**
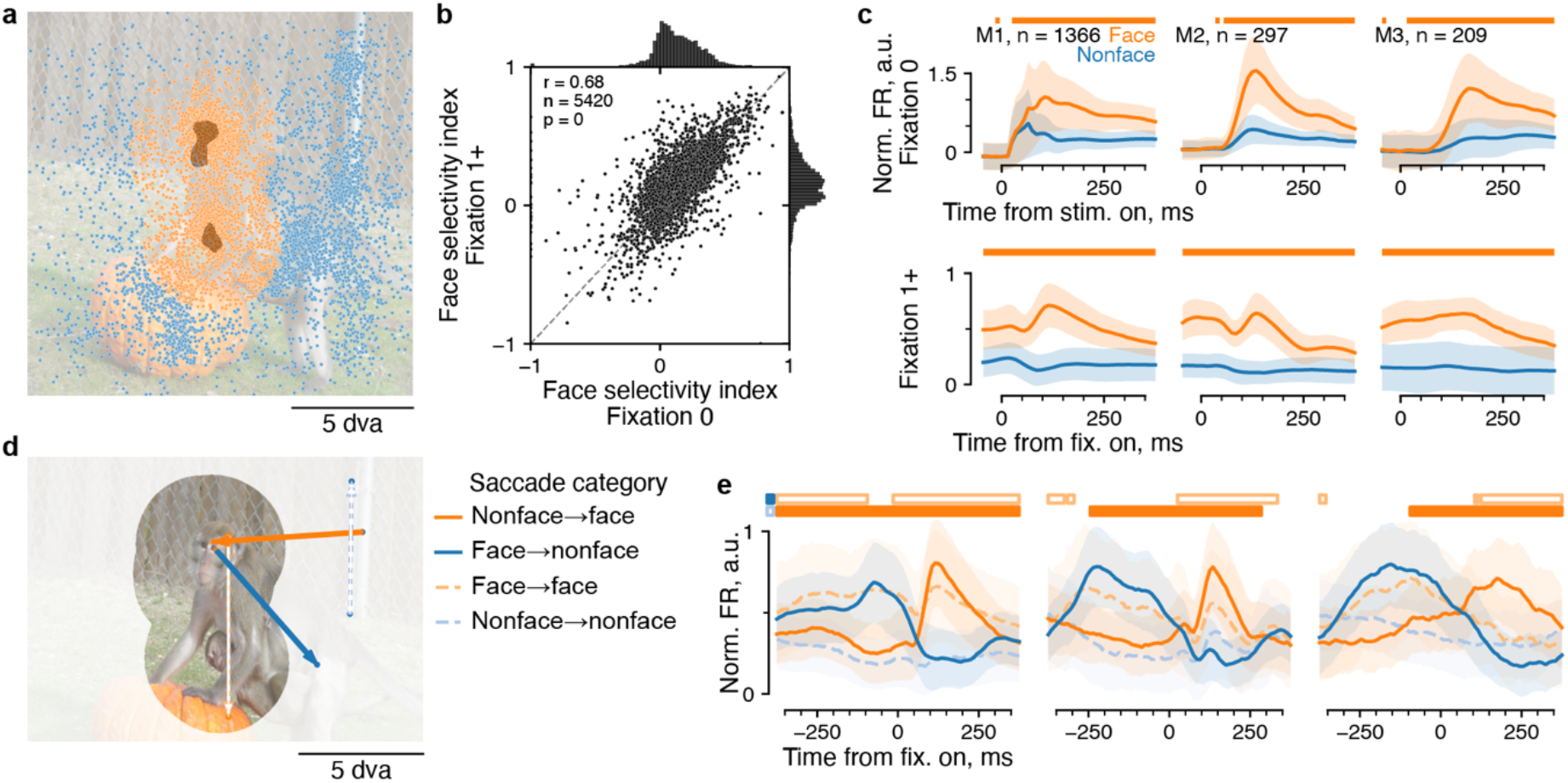
Face-selective neurons responded to whether fixations landed near a face. **a**, Face ROIs were used to categorize fixations, illustrated here for the same example image as in Fig. 1b. The dark shaded regions correspond to face ROIs; each dot indicates a fixation; color indicates the categorization of the fixation content (orange, face; blue, nonface). **b**, Face selectivity was quantified by a selectivity index (Methods) and compared between the zeroth (x-axis) and subsequent fixations (y-axis). The dashed line is the identity line. **c**, Mean responses per category aligned to the onset of either the zeroth (top row) or subsequent (bottom row) fixations. Each column corresponds to one monkey; n indicates the number of neurons; lines and shading indicate mean ± stdev. across neurons; horizontal bars indicate time bins where responses were significantly greater for face than nonface responses (p < 0.01, one-tailed permutation test, FDR-corrected). **d**, Non-zeroth fixations were further divided according to their preceding fixations into two-by-two categories. An example saccade is shown for each category. Dashed arrows indicate saccades between the same fixation category (blue: nonface-to-nonface; orange: face-to-face). Solid arrows indicate saccades between different categories (blue: face-to-nonface; orange: nonface-to-face). **e**, Mean responses for each of the saccade categories shown in **d** for the same neurons as in **c**. Each subplot shows data from one monkey. Horizontal bars indicate time bins where responses were significantly greater for nonface-to-face than for nonface-to-nonface saccades (lower solid bars), or face-to-face than for face-to-nonface saccades (upper open bars). Other plotting conventions follow those in **c**.

During active vision, face-selective responses began before fixation onset (Fig. 2c, bottom row). However, because face fixations often preceded other face fixations, the apparent predictive selectivity could be due to the previous fixation. To test this possibility, we split neighboring fixations into four conditions by the categories before and after the current saccade (Fig. 2d). We further restricted our analysis to large saccades (> 4 dva) to accommodate the imprecision of arbitrary ROI boundaries. Face neuron responses meticulously followed the category of each fixation across saccades (Fig. 2e). For instance, across nonface-to-face saccades, face neuron firing rates switched from low to high within tens of milliseconds after fixation onset. While the fixation category explained broad trends in the response magnitude, it also revealed fine differences. For example, face responses following another face fixation were lower than face responses following a nonface fixation (Fig. 2e; compare dashed orange lines to solid lines in the peak after fixation onset). Moreover, when comparing nonface-to-face saccades with nonface-to-nonface saccades, responses were significantly higher well before saccade onset in all monkeys (Fig. 2e). We could not rule out the possibility that this presaccadic difference was due to our conservative ROIs, or because the 5-dva RF size we assumed did not fully account for the tapering periphery of RFs. Therefore, we looked for an analysis that depended less on a precise, binary RF delineation and sought a metric applicable to neurons without a subjectively defined selectivity.

### Ventral visual responses were specific to individual fixations

The selectivity of ventral visual neurons did not always correspond to clear categories; even if it did, it would be problematic to interpret ROIs assuming binary selectivity. Instead, we used return fixations to define a self-consistency metric. Primates repeatedly foveate the same location above chance frequency in various behavioral contexts, including visual search and free viewing (Zhang et al., 2022). Fig. 3a shows example return fixations pairs (black lines) in one example session, within and between trials. Return fixation pairs should lead to similar responses if neurons encode the retinotopic stimulus content at each fixation. We quantified the self-consistency of responses during return fixations (Fig. 3b) by calculating Pearson’s correlation coefficient of responses between every pair of return fixations, across different return fixation pairs. This quantification is analogous to how studies typically quantify response self-consistency as the correlation between repeated presentations across stimuli. To assess temporal specificity, we calculated self-consistency for responses at different times relative to fixation onset. The first two subplots in Fig. 3c, corresponding to the purple bars in Fig. 3b, show the firing rates of an example neuron 200 ms before (subplot 1) and after (subplot 2) the onset of return fixations. The responses were more consistent after (r = 0.55) than before (r = 0.30) fixation onset.

**Fig. 3.**
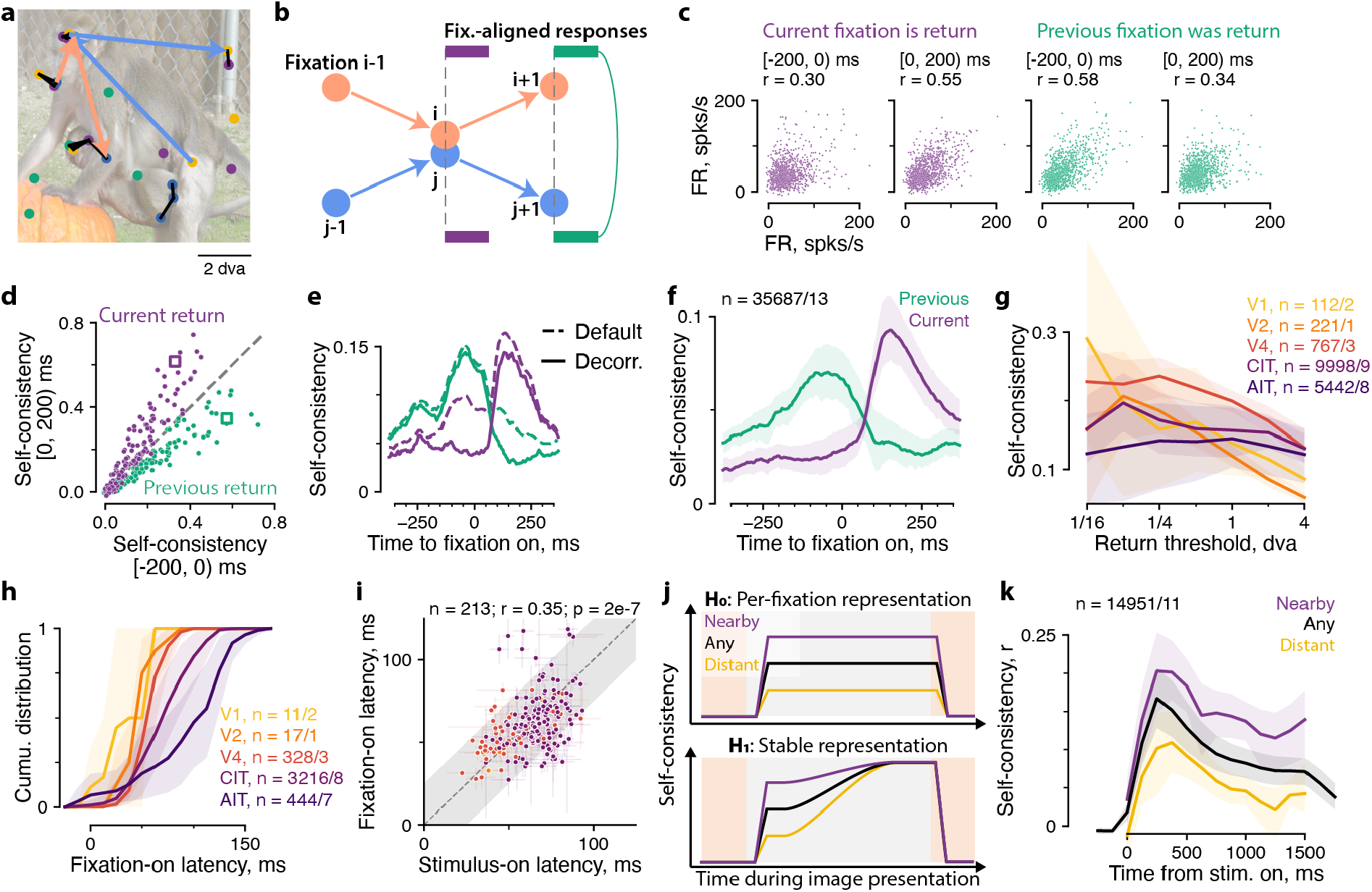
Response self-consistency indicated specificity to each fixation but no stable representation across fixations. **a**, Example return fixation pairs. A return fixation pair is two nearby (< 1 dva) fixations on an image, within a trial or across trials. The dots indicate fixations, color-coded by trial. Black lines join return fixation pairs. **b**, The schematic illustrates two pairing rules for calculating response self-consistency based on return fixations. Orange and blue indicate two fixation sequences. The second fixation in each sequence (denoted i and j, respectively) make up a return pair. Neuronal responses aligned to the second fixation onset (purple) are paired based on the rule, ‘the current fixation is a return fixation.’ Responses aligned to the third fixation onset (green) are paired based on the rule, ‘the previous fixation was a return fixation.’ **c–e**, Example data illustrate how we quantified self-consistency. **c**, Each dot indicates the firing rates (FR) of an example neuron in a return fixation pair; the x- and y-axis show responses to one or the other fixation in each pair. The four subplots correspond to two response time bins: 200 ms before (columns 1 and 3) or after (columns 2 and 4) fixation onset; and two response pairing rules: paired by the current (columns 1 and 2) or previous (columns 3 and 4) fixation. We quantified self-consistency as Pearson’s correlation of paired firing rates across pairs. A small amount of random noise was added for visualization purposes only because firing rates were discrete and often overlapped. **d**, Self-consistency for all neurons in this example session. The x- and y-axes correspond to the two response time bins. The colors indicate the response pairing rule (previous or current; see **b, c**). Within a color, each dot indicates one neuron. Square markers indicate the example neuron in **c**. The dashed line is the identity line. **e**, Self-consistency time courses were calculated using 50-ms sliding response time bins. The lines indicate averages over neurons in the example session. Dashed lines correspond to all return pairs (‘default’); solid lines correspond to ‘decorrelated’ return pairs (see text). **f**, Decorrelated self-consistency time courses summarized over monkeys and neurons. **g**, Self-consistency as a function of the threshold for pairing return fixations, separately for visual areas. **h**, Cumulative distribution of response latency following fixation onset, separately for visual areas. **i**, Comparison of response latency following stimulus onset (x-axis) and fixation onset (y-axis). Each dot indicates one neuron; error bars indicate bootstrap stdev. of the latency estimates; color indicates visual area as in **h**. Grey diagonal shading indicates 25 ms, the threshold bootstrap stdev. of latency estimates. **j**, Schematics of prediction by the null hypothesis of purely retinotopic responses (H_0_, top) and the alternative hypotheses of stable representation (H_1_, bottom; see text). Colors indicate various measures of response self-consistency. **k**, Mean self-consistency as a function of time from stimulus onset, to compare against the predictions in **j**. In **f, g, h**, and **k**, lines and shading indicate hierarchical mean ± bootstrap 95%-CI, first over neurons, then over monkeys. The n values indicate the total number of neurons/monkeys.

For this example neuron, self-consistency was well above zero before fixation onset. We realized that fixations close by in time also tended to be close by in space. To account for the spatial correlation between subsequent fixations, we contrasted two pairing rules for comparing responses: pairing responses such that either the *current* fixations (response times t ≥ 0) or the *previous* ones (t < 0) were return fixations (respectively purple and green in Fig. 3b). The four subplots in Fig. 3c illustrate response self-consistency for the example neuron at different response windows for different paring rules. While the responses were similar even before fixation onset (Fig. 3c, first subplot), this similarity was better explained by the previous fixation because the responses were more self-consistent when the previous fixations were paired (Fig. 3c, third subplot) than when the current ones were. The converse was true for responses after fixation onset (Fig. 3c, second and fourth subplots).

Most neurons in this example session showed higher self-consistency in the response time window corresponding to the paired fixation than in the complementary window (Fig. 3d). We can reveal this temporal specificity in higher resolution by using 50-ms sliding time bins (Fig. 3e, dashed lines). Consistent with the example neuron in Fig. 3c and the temporally coarser results in Fig. 3d, the dashed lines in Fig. 3e show that neuronal responses followed the contents of each fixation. When the current fixation was a return fixation (purple), self-consistency was low before fixation onset and increased after it. Conversely, when the previous fixation was a return fixation (green), self-consistency was high before the (current) fixation onset and decreased following it.

We devised a further control for the above-zero self-consistency during the non-paired period by decorrelating the non-paired fixation. For each pairing rule (previous or current), we sub-selected fixation pairs such that fixations during the non-paired period were more than 4 dva apart. The decorrelation procedure reduced self-consistency in the non-paired period (solid lines in Fig. 3e) and barely affected self-consistency values within the paired period. Fig. 3f summarizes the time course for the decorrelated self-consistency, showing robust specificity to each fixation across monkeys and neurons. We used the decorrelated variant of self-consistency in further analyses.

Return fixation self-consistency furnished a measure for the spatial specificity of neuronal responses during free viewing. We asked how self-consistency depended on the distance cutoff for defining return fixations. Self-consistency increased with smaller, more stringent thresholds, plateauing at 1 dva for anterior IT (AIT) neurons and continuing to increase in central IT (CIT), V4, V2, and V1 down to 1/4 dva (Fig. 3g), the resolution of our eye trackers. Thus, ventral visual responses were retinotopically precise during free viewing of dense natural scenes.

The time course of self-consistency also supplied a measure of response latency during free viewing. We estimated the latency of fixation-specific responses as the time point at which responses became better explained by the current than the previous fixation, i.e., the crossing point between the previous and current return self-consistency curves. This latency estimate yielded typical values that, as expected, increased along the processing hierarchy in the ventral pathway (Fig. 3h). We could further compare, in the same neurons, the fixation-onset latency and the classical, stimulus-onset latency, using zeroth fixation responses to estimate the latter (see Methods). For a small subset of neurons, we could estimate both metrics reliably (bootstrap stdev. < 25 ms; Methods). The two response latency measures were similar and showed consistent differences across neurons (Fig. 3i).

The results thus far indicate that ventral visual responses tracked individual fixations during free viewing. The possibility remains that some subtle aspect of the responses may integrate across fixations to build up a stable representation. We hypothesized that a stable representation would manifest as reduced retinotopic specificity over multiple fixations on the same image. Under the null hypothesis that responses remain retinotopic (Fig. 3j top, H_0_), there should be a persistent gap throughout the trial between different measures of response similarity: self-consistency between return fixations < 1 dva apart (purple); response similarity between any two fixations on the same image (black); and response similarity between distant fixations > 8 dva apart (orange). Return fixations had similar retinal stimulation and should correspond to higher self-consistency, whereas distant fixations had dissimilar retinal stimulation and should correspond to low response similarity. Response similarity should be intermediate between two random fixations on the same image. In contrast, if retinotopic specificity decreased throughout continued viewing of the same image, the gap should narrow, if not vanish, between the three response similarity measures (Fig. 3j, bottom, alternative hypothesis H_1_), meaning that any fixation location led to a similar response. We evaluated the three measures of response similarity (i.e., self-consistency) and summarized them across all trials lasting 1.5 s (Fig. 3k; Methods). A constant retinotopic gap persisted among the three self-consistency measures, consistent with the null hypothesis of retinotopy and contradicting the alternative hypothesis of a steadily accumulating stable representation. However, the null hypothesis did not account for a general drop in self-consistency throughout the trial. We speculate that this drop is due to the general decrease in firing rate as the monkeys continued to view the same image (Fig. 1g), consistent with adaptation.

In sum, the spatiotemporal properties of response self-consistency indicated that, during active viewing, ventral visual neurons remained retinotopically specific and showed response dynamics governed by the contents of the current fixation. Neurons maintained their retinotopic specificity during continued viewing of an image over 1.5 s without integrating multiple fixations to reduce specificity to each gaze.

### Computational models predicted ventral visual responses from fixation-aligned stimulus content

The response properties described so far encouraged us to test whether image-computable models could predict neuronal activity during free viewing. Previous work showed that artificial neural networks provided a reasonable approximation to visual responses obtained during passive viewing (Schrimpf et al., 2018; Yamins et al., 2014). We adapted this class of models to free viewing responses. We fit the models to predict fixation-aligned responses from a 4 × 4 dva image patch (i.e., 1/16^th^ the area of a 16 × 16 dva stimulus) centered on each fixation (Fig. 4a). A pretrained artificial neural network (vision transformer) (Dosovitskiy et al., 2020) converted each image patch into a 1024-D feature vector. We fit a linear model to predict neuronal responses from the feature vectors and evaluated model performance using cross-validation across images. While previous studies have shown similar models to capture trial-averaged responses during controlled fixation, we found the models could also predict neuronal responses during individual fixations. The models captured a large fraction of the explainable (i.e., self-consistent) responses during both the passive viewing-like zeroth fixation (Fig. 4b, Extended Data Fig. 1a) and free viewing (Fig. 4c, Extended Data Fig. 1b). The fraction of responses explained, which normalizes model performance by the self-consistency ceiling, peaked at 52% during free viewing (95%-CI: 43%–61%; Extended Data Fig. 1), in line with the performance of related models in passive viewing studies (Cadena et al., 2019; Schrimpf et al., 2018; Yamins et al., 2014).

**Fig. 4.**
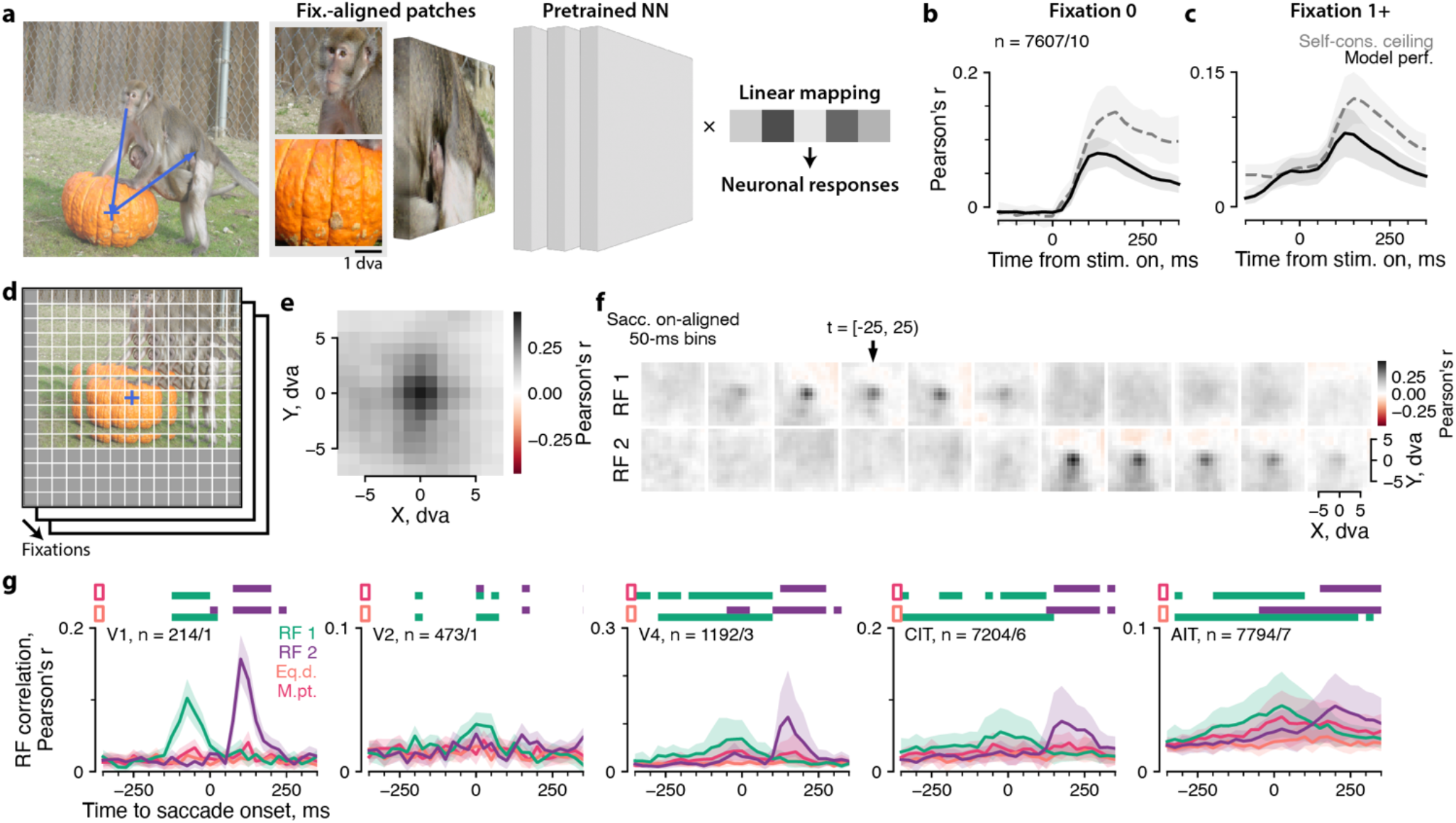
Computational models captured neuronal selectivity to individual fixations and revealed the spatiotemporal structure of receptive fields. **a**, We adapted computational models of visual neuronal responses to single-fixation free-viewing data. The model comprised a pretrained, fixed neural network (NN) feature extractor (vision transformer; see Methods for details) and a linear mapping fit to neuronal responses. Model inputs were fixation-centered image patches 4 dva in size (illustrated in **a**) in panels **b** and **c**, or 2 dva in panels **d–g. b, c**, Cross-validated model performance compared to the ceiling of return fixation self-consistency, separately for fixation 0, **b**, or subsequent fixations 1+, **c. d**, Illustration of model-based inference of neuronal receptive fields (RFs) during free viewing. We partitioned the retinotopic image into 2-dva patches on a grid of offsets centered on the fixation indicated by a cross (also in **a**). Other fixations were also partitioned with the same grid. The model NN converted each patch to a feature vector. Then, we evaluated the feature fit to neuronal responses at every offset from fixation (see the main text). This process derived an RF map consisted of model-based, cross-validated local stimulus ‘correlation’ to neuronal responses across fixations, quantified by Pearson’s r. **e**, RF inferred from the zeroth-fixation responses of an example neuron. **f**, Spatiotemporal RFs inferred for the example neuron using responses 50-ms rolling time bins aligned to saccade onset. The arrow indicates the bin centered on saccade onset. The two rows correspond to RFs centered on the saccade starting (RF 1, top) or ending point (RF 2, bottom). **g**, Quantification of RF presence over time and coordinate frames. Colors indicate RF 1 (green), RF 2 (purple), the equidistant control (peach), and the midpoint control (magenta; see the main text). Each subplot corresponds to a visual area. Horizontal bars indicate statistically significant differences from the equidistant (lower two rows) or midpoint control (upper two rows; p < 0.01, one-tailed permutation test, FDR-corrected). In **b** and **g**, lines and shading indicate hierarchical mean ± bootstrap 95%-CI, first over neurons, then over monkeys. The n values indicate the total number of neurons/monkeys.

### Stimulus-selectivity implied retinotopic receptive fields that shifted with each fixation

Insofar as the computational models could capture neuronal stimulus selectivity, they provided a means to infer the structure of neuronal receptive fields (RFs) during natural vision. The RF corresponds to a spatiotemporal section in the visual input that explains a neuron’s responses. To localize this explanatory section, we reasoned that stimulus contents within the RF should allow a model to predict neuronal responses, while stimulus contents outside the RF should not allow for significant prediction. Therefore, we partitioned the visual scene centered on each fixation into a grid of small, overlapping image patches 2 × 2 dva in size and separated by 1-dva intervals (Fig. 4d). A separate model instance used the patch at each offset from fixation to predict neuronal responses. The resulting map of model performance at different offsets would reveal the spatial extent of a neuron’s RF. Empirically, we found it helpful to further regularize this process by sharing model coefficients across offset locations (see Methods for details), resulting in a metric reminiscent of reverse correlation. Using simulated responses, we validated that this model-based, cross-validated reverse correlation procedure for inferring RF structure could recover the location, size, and shape of ground truth RFs that generated simulated responses (Extended Data Fig. 2).

Fig. 4e shows the RF inferred from responses 70–200 ms following zeroth fixations for an example CIT neuron. The map showed a focal region of high correlation; its location and extent were consistent with results from conventional mapping experiments in this chronic recording array (Extended Data Fig. 3, Pa array 1). The mapping method recovered zeroth-fixation RFs that matched conventionally mapped RFs across recording arrays (Extended Data Fig. 3).

The model-based RF mapping procedure allowed us to examine the dynamics of RF remapping directly during free-viewing behavior with naturalistic stimuli. We evaluated RFs using rolling response time bins aligned to saccade onset, anchoring the RF alternatively on the fixation point before or after the saccade (FP 1 or FP 2). To help disambiguate the RF in the two coordinate frames, RF 1 or RF 2, we included only large saccades (> 5 dva) in this analysis. Fig. 4f shows the two sets of inferred spatiotemporal RFs for the same example neuron in Fig. 4e. When an RF was present, it was focal in one of the two fixation-centered coordinate frames. The RF shifted from FP 1 to FP 2 in the response time bin 75–125 ms following saccade onset, consistent with the typical latency in central IT and an average saccade duration of around 50 ms.

To summarize the time course of RF shifts across sessions, we quantified RF presence using its consistency across cross-validation splits, regularized via 2D Gaussian fits (Fig 4g; Methods). We quantified each time bin and RF coordinate frame independently, to allow for any potential shifts in the RF that could be non-parallel to the saccade (Neupane et al., 2016; Tolias et al., 2001; Zirnsak et al., 2014). The conclusions remained similar if we quantified perisaccadic RFs by their consistency to the zeroth fixation RF (Extended Data Fig. 4a). Across visual areas, RF presence was strongest in the period corresponding to the anchoring fixation, i.e., before the saccade for RF 1 (green in Fig. 4g) and after the saccade for RF 2 (purple in Fig. 4g). The strength of evidence for the two RFs crossed over at times that were broadly consistent with neuronal response latencies that increased along the processing hierarchy (from left to right in Fig. 4g).

In some visual areas, there was some above-zero RF2 evidence before the saccade (for example, in V4 and AIT for RF 2, purple), reminiscent of the above-zero baseline in the self-consistency time course (Fig. 3). This subtle but quantifiable RF evidence in the non-anchoring fixation could indicate predictive RF remapping (purple, t < 0) or, alternatively, reflect stimulus feature correlations that the saccade size criterion did not eliminate. To distinguish between these two possibilities, we included two control analyses by inferring RFs anchored on two types of counterfactual points: a point equidistant to FPs 1 and 2 but unvisited by the eye or the midpoint between FPs 1 and 2 that the eye quickly passed over. The equidistant control should capture (isotropic) spatial autocorrelations in the stimulus. The midpoint control should capture any stimulus features shared between FPs 1 and 2. The evidence for RF 2 significantly exceeded the evidence for the counterfactual equidistant RF before saccade in V1, V4, and AIT (Fig. 4g, lower horizontal bars; p < 0.01, one-tailed permutation tests, FDR-corrected).

However, no RF 2 evidence before saccade onset exceeded the evidence for the counterfactual midpoint control or, indeed, RF 1 evidence in any visual area. The early RF 2s also looked weaker than retinotopic RFs for example arrays that showed statistical significance for prediction and were not always consistent with zeroth-fixation RFs (Extended Data Fig. 5; first row in each group). We further tested whether the putative predictive remapping was attributable to fixation 1 by varying its onset time. If the early RF 2 evidence was predictive, we expected it to align to saccade onset rather than fixation 1 onset. Instead, the early RF 2 evidence, when it remained, shifted earlier for earlier fixation 1 onsets (Extended Data Fig. 5, comparing second and third rows in each group). Thus, while we cannot rule out predictive remapping, the results suggest that any putative remapping factored less in ventral visual responses than the current RF during free viewing.

## Discussion

We studied neurons along the ventral visual cortex, one of two major cortical visual pathways in primates, while monkeys freely viewed natural images. We aimed to design robust analyses that could uncover the general principles governing visual responses during this unconstrained behavior. We found that neurons in the ventral visual pathway maintained their feature selectivity, yoked their responses to eye movements, and responded with classical latencies following fixation shifts. When monkeys viewed images containing faces, face neurons responded more strongly when the monkey fixated near faces than further away, and the responses refreshed after each gaze shift (Fig. 2). Neurons responded consistently during fixations on similar locations (Fig. 3a–d). This consistency evinced high spatial precision (Fig. 3g) and classical response latencies (Fig. 3e, f) that increased along the ventral processing hierarchy (Fig. 3h, i). Throughout the 1.5 seconds an image remained on screen, neuronal responses remained tightly linked to the present fixation and did not become invariant to gaze (Fig. 3j, k). Synthesizing these principles in computational models, we could explain a large fraction of stimulus-selective neuronal responses in single fixations (Fig. 4a, b), extending work from passive viewing studies (Yamins et al., 2014). Using these models as a pivot, we factorized out stimulus selectivity to reveal the spatiotemporal structure of neuronal receptive fields (RFs) during free viewing (Fig. 4c–g). The inferred RFs corroborated response self-consistency to show that ventral neurons responded to a spatially and temporally local part of the retinotopic stimulus.

We developed several analysis methods that may benefit future studies involving unconstrained visual behaviors. The return self-consistency metric quantifies the response variation explained by controlled visual inputs and reveals the spatial specificity and temporal dynamics of the visual responses. Furthermore, we showed that image-computable models can capture the stimulus selectivity in single-trial free viewing responses. Finally, we developed a model-based method to infer RFs during natural vision, extending existing methods that depend on idealized tuning functions or ad hoc feature maps (Kay et al., 2013; Yates et al., 2021).

Ventral visual neurons have been traditionally understood as retinotopic feature detectors. Much of this knowledge derives from experiments that deliberately limit spatial and temporal context and exclude eye movements. Some studies using more natural viewing conditions hint at non-classical responses (Gallant et al., 1998; McMahon et al., 2015; Podvalny et al., 2017; Rolls et al., 2003; Russ et al., 2023; Sheinberg & Logothetis, 2001) but do not identify the specific mechanisms that could have led to nonretinotopic responses. Others suggest that neurons in V1 and IT maintain their classical response properties during free viewing (DiCarlo & Maunsell, 2000; Livingstone et al., 1996). We collected an extensive dataset that allowed for detailed analyses and computational modeling. Our results support the view that ventral visual responses were predominantly retinotopic during natural vision. At the same time, we found two deviations from strictly classical responses. First, firing rates and selectivity generally decreased during the trial (Figs. 1g, 3k). Future work is needed to evaluate whether this may be attributable to response adaptation mechanisms, potentially at the network level (Solomon & Kohn, 2014; Vinken et al., 2020). Second, we could not completely rule out predictive remapping (Fig. 4g), which we discuss in more detail below.

We closely examined predictive remapping as a candidate non-classical response property. Predictive remapping refers to the anticipatory updating of retinotopic receptive fields before the eye moves. Studies have reported predictive RFs primarily in dorsal cortical and subcortical areas such as the lateral intraparietal area (Duhamel et al., 1992), frontal eye field (Umeno & Goldberg, 1997; Zirnsak et al., 2014), superior colliculus (Walker et al., 1995), area MST (Inaba & Kawano, 2014), and area V3A (Nakamura & Colby, 2002). By comparison, evidence for predictive remapping is limited in the ventral visual cortex (Neupane et al., 2016; Tolias et al., 2001). We found statistical evidence that neurons might predictively remap in V4 and anterior IT (Fig. 4g) when compared to an equidistant condition that controls for natural image statistics assuming isotropic autocorrelation. However, we could not rule out all potential confounds. The equidistant point is on average closer to the image boundary, potentially causing large RFs to be truncated more often (Extended Data Fig. 4b). Meanwhile, the monkey may prefer salient parts of the image that were more correlated with each other than the equidistant point, violating the isotropic assumption. This may underlie the spurious evidence for prediction in simulated responses for small RFs (Extended Data Fig. 2, first row). Finally, it bears emphasis that even if the statistical difference reflected actual neuronal mechanisms, the evidence for the current RF overshadows that for the putative predictive RF.

Our study also differs in multiple key aspects from V4 studies that report remapping. Tolias et al. (2001) used a persistent, cell-specific stimulus such as an oriented bar. Neupane et al. (2016) used a flashed probe—a square light spot on a black background. Both studies showed the probe on an otherwise empty display and varied the stimulus location but not the identity. We studied vision in dense, persistent visual scenes that continuously stimulated neurons in their classical RFs. Thus, we quantified how neuronal responses corresponded to stimulus content instead of evaluating the presence or absence of responses as a function of stimulus location. It is unknown whether previously reported predictive RFs show the same feature selectivity as classical RFs. Alternatively, shifting RFs could solely indicate the presence of a salient stimulus without its content, consistent with the interpretation of predictive RFs as attention remapping (Rolfs, 2015). Attentional remapping could explain why superior collicular remapping diminishes when probed with multiple simultaneous stimuli (Churan et al., 2011), a stimulation condition that is in turn closer to free viewing of dense natural scenes. The potential interplay among attention, selectivity, and stimulation conditions suggests caution when extrapolating from highly constrained experiments to explaining vision during natural behavior.

Indeed, there is no reason to expect that seeing involves a detailed, stable map of the visual scene. Conscious perception probably does not involve a veridical image of the world; idioms like ‘out of sight, out of mind’ give voice to this near-truism. Empirical support for the absence of perfect visual stability comes from findings of change or inattentional blindness (Mack & Rock, 1998), memoryless visual search (Horowitz & Wolfe, 1998; Zhang et al., 2018), and perisaccadic mislocalization (Matin & Pearce, 1965; Ross et al., 2001). Across saccades, people can miss even relatively large object displacements (Bridgeman et al., 1975). However, behaviors like the multi-step saccade task (Hallett & Lightstone, 1976) require the subject to maintain some information across eye movements. While the nature and content of stable visual perception are unclear, existing evidence suggests that the brain may not stabilize the rich representations of ventral vision but maintains only the information necessary for guiding specific actions.

Finally, we underscore the value of testing theories of brain function under natural behavior. Reductionist experiments can illuminate the mechanisms of cognition only as long as the isolated facets reflect how the brain operates during natural conditions. The brain evolved for behavior, with which neuroscience should start and end (Krakauer et al., 2017; Leopold & Park, 2020). Natural behavior is a source for generating hypotheses and should be the final test for principles gleaned from artificial experiments. We have developed flexible analyses that can be applied to unravel visual response properties across brain areas during other natural behaviors.

## Methods

### Experiment details

#### Subjects

All procedures were approved by the Harvard Medical School Institutional Animal Care and Use Committee and conformed to NIH guidelines provided in the Guide for the Care and Use of Laboratory Animals. Eleven adult *Macaca mulatta* (one female, ten males; 5–13 kg; 2–17 years old) and two adult male *Macaca nemestrina* (13 and 15 kg; 12 and 14 years old) were socially housed in standard quad cages on 12/12 hour light/dark cycles.

#### Surgical procedures

Animals were implanted with custom-made titanium or plastic headposts before fixation training. After several weeks of fixation training, the animals underwent secondary surgeries for array implantation. All surgeries were done under full surgical anesthesia using sterile technique.

#### Physiological recording

Animals were implanted with custom floating microelectrode arrays (32 channels, MicroProbes, Gaithersburg, MD or 128 channels, NeuroNexus, Ann Arbor, MI) or microwire bundles (64 channels; MicroProbes). Each animal received 1–5 arrays throughout data collection spanning three years. Neural signals were amplified and sampled at 40 kHz using a data acquisition system (OmniPlex, Plexon, Dallas, TX). Multi-unit spiking activity was detected using a threshold-crossing criterion. Channels containing separable waveforms were sorted online using a template-matching algorithm.

#### Behavioral task

Monkeys performed a free viewing task with a range of parameters. Images were typically presented at a size of 16 × 16 degrees of visual angle, but some experiments used other sizes ranging from 8 × 8 to 26 × 26 dva. Most experiments used a 1.5 s presentation duration, but some experiments used other durations ranging from 0.3 to 60 s. Image presentation was pseudorandomly ordered in a block design such that images were repeated when all images had been shown once. Image position was randomly shifted each presentation to encourage free looking because most monkeys have been extensively trained to fixate. Monkeys were rewarded at random intervals with a drop of juice for maintaining their gaze within a window around the image. Task control was handled by a MATLAB-based toolbox, NIMH MonkeyLogic (Hwang et al., 2019). The task-control software monitored and recorded eye position from an infrared eye tracker (ISCAN, Woburn, MA, or EyeLink, Ottawa, Canada). Eye tracking was calibrated before each session using a projective transform.

### Quantification and statistical analysis

#### Data preprocessing

The neural recording was synchronized by TTL events to task control and behavior data including eye tracking. Stimulus onset times were refined using a photodiode signal. We measured and corrected for a fixed latency per tracker in the eye signal. Using a motorized model eye attached to a potentiometer, we compared the eye position signal recorded as a voltage trace alongside neural data to the synchronized behavior data stream to obtain a constant offset. This constant offset was applied before downstream analysis. Fixations and saccades were detected using ClusterFix (König & Buffalo, 2014) with default parameters.

#### Neuron selection

Because we recorded from chronic multi-electrode arrays, not all channels contained visually responsive signals. We elected to include only visually selective units in all analyses. Operationally, we defined visually selective units as those that passed a threshold of r = 0.1 on return fixation self-consistency with a 250-ms response window based on either previous or current return; see the section, *Self-consistency metrics* below for how this metric was defined. This criterion included 48 ± 23% of units (mean ± stdev.).

#### Fixation selection

Fixations were included in analyses based on the following criteria: 1) the fixation lasted at least 100 ms; 2) the fixation landed within the image.

#### Face-specific analysis (Fig. 2)

Face regions of interest were either manually drawn for datasets containing both monkey and human faces or detected as bounding boxes using a pre-trained face-detection neural network (RetinaFace; Deng et al. (2020)) for datasets containing human faces only. Fixations were classified as face fixations if they landed within 5 dva of a face ROI and nonface otherwise. To match the face and nonface conditions more closely, we only considered nonface fixations on images that contained a face ROI. Face-selectivity index (FSI) was calculated using the responses 70–200 ms from stimulus onset (‘fixation 0’) or fixation onset (‘fixation 1+’) as (a – b) / (a + b), where a and b corresponds to face and nonface fixation responses, respectively. We restricted the face-specific analysis to three monkeys with face-patch arrays and only those units with an FSI > 1/3 based on fixation 0 responses. Responses were normalized per neuron to the 95^th^ percentile across fixation types (0 or 1+), categories (face or nonface), and time points.

#### Return fixation self-consistency (Fig. 3)

In standard controlled viewing experiments, self-consistency is typically defined using the split-half correlation of trial-averaged responses conditioned on stimulus identity. We generalized this metric to free viewing. First, we relaxed trial-averaging by considering ‘single-trial’, i.e., single-fixation, responses. Second, we considered return fixations to be analogous to repeated presentations. In the ‘default’ setting, all fixations on the same image in each session were paired based on a proximity threshold, typically 1 dva. Two adjacent fixations tended to be close by (Fig. 1e), leading to nearby locations in the non-paired fixation. To reduce this correlation in the non-paired fixation (i.e., the preceding fixation when the ‘current’ fixation was paired and the subsequent fixation when the ‘previous’ fixation was paired), we further sub-selected return fixation pairs where the non-paired fixation was separated by > 4 dva. Neuronal responses were aligned to fixation onset and paired based on the corresponding fixations. Responses were calculated using either the 250 ms before and after fixation onset or using rolling 50-ms time bins with 25-ms steps. Self-consistency was quantified by Pearson’s r.

In Fig. 3k, we computed variants of self-consistency conditioned on different notions of repeat trials. ‘Same image’ self-consistency compared all fixation pairs within the same image, across images, regardless of fixation location. ‘Distant fixation’ self-consistency compared all fixation pairs that were > 8 dva apart. In this analysis, responses were aligned to stimulus rather than fixation onset and computed in 250-ms time bins with 125-ms steps. Every fixation was assigned by its onset time to a time bin; a time bin may contain multiple fixations. Two time bins were compared if they contained any fixations that matched a fixation pairing rule.

#### Estimates of response latency (Fig. 3h–i)

Fixation response latency was estimated as the first time point on either side of time 0 that the ‘current’ return fixation self-consistency exceeded the ‘previous,’ both using decorrelated return fixation pairs. To estimate stimulus-onset response latency, we elected to use self-consistency instead of the traditional average firing rate to be more comparable to the fixation-onset latency. We calculated return fixation self-consistency using zeroth fixations exclusively. Response latency was estimated as the closest time point on either side of time 0 that the self-consistency time course crossed from below to above a threshold, which was in turn based on the average of the 2.5^th^ and 97.5^th^ percentiles of the self-consistency time course. For all self-consistency metrics, bootstrap estimates were obtained by sampling fixation pairs with replacement. We estimated latency separately for each self-consistency bootstrap sample to obtain a bootstrap distribution of latency estimates. In Fig. 3k, we included only units that had both well-estimated latencies based on several quality control criteria: 1) bootstrap stdev. < 25 ms; 2) bootstrap bias < 12.5 ms; 3) peak self-consistency > 0.1; 4) crossing point was unique for 100 ms on either side.

#### Computational model of neuronal responses (Fig. 4a–b)

Image patches were embedded in the feature space of a neural network (NN), a pre-trained Vision Transformer (ViT) (Dosovitskiy et al., 2020). We used the model instance, ‘vit_large_patch16_384’ in the Python library ‘pytorch image models’ (Wightman, 2019) and extracted features from the layer, ‘blocks.10.norm1’. The features were average over the sequence dimension to result in a 1024-dimensional vector. To efficiently fit the linear mapping models, we pre-calculated and cached the NN representations using a discrete sampling grid, either 4 × 4 dva patches in 1-dva steps for fixation-centered models (Fig. 4a–b), or 2 × 2 dva patches in 0.5-dva steps for inferring receptive fields (Fig. 4c–f). Patches extending beyond the image extent were padded with gray. Each fixation was digitized into the closest bin to obtain the feature vector of the corresponding image patch. Responses were calculated per fixation using a 50-ms window aligned to fixation onset. A Ridge regression model was fit to map between fixation-aligned model features and neuronal responses, with the regularization parameter alpha = 5500. The linear mappings were fitted and evaluated using five-fold cross-validation across images. Thus, no return fixations were present in both the training and testing sets. Model performance was quantified by the correlation (Pearson’s r) between predicted and actual responses on held-out fixations. Ceiling-normalized model performance was calculated by dividing model performance with return fixation self-consistency, clipping the result between 0 and 1, then squaring it. This measure would correspond to the fraction of variance explained up to an optimal linear transformation.

#### Model-based inference of receptive field structure (Fig. 4c–f)

At each fixation, a grid of 2 × 2 dva image patches was extracted on a fixation-anchored grid of offset locations, from -7 to 7 dva in 1-dva steps. Each image patch was converted into a 1024-D model embedding vector, resulting in a 15 × 15 × 1024 retinotopic stimulus representation akin to a multichannel image. At each of the 15 × 15 = 225 offset locations, a separate linear mapping was fit and evaluated across fixations and cross-validated over images as described above. This process resulted in a 15 × 15 × 5 map of model performance per cross-validation (CV) split.

To further regularize this map, we took the 1024-D model weights from the location of peak performance per CV split. The model weights were applied to held-out fixations to project each 15 × 15 × 1024 fixation-centered stimulus representation to a 15 × 15 scalar map, akin to a grayscale image, that was private to each neuron, response window, and CV split. These scalar maps were correlated to fixation-aligned neuronal responses, a process analogous to reverse correlation. The correlation was performed either across CV splits to obtain a single map per condition, or within each split for later comparison across splits as described below. Maps were clipped at 0 because negative correlation indicated over-fitting and squared because doing so resulted in better Gaussian fits (described below).

To quantify the clear and consistent presence of receptive fields (RFs), we fitted a Gaussian distribution to the inferred RF per CV split, then evaluated the goodness-of-fit on the 15 × 15 RF map from other splits. Goodness-of-fit was quantified with Pearson’s r and averaged over 5 × 4 pairs of splits (the pairs were directional because only one split contributed to the Gaussian fit).

The process above inferred the spatial structure of an RF and quantified its clear presence separately for each neuron and response time window. This process was repeated over neurons, separately per response window to allow for potential changes in stimulus selectivity and RF structure across time. In Fig. 4d, a fixational RF was inferred for the response window 70–200 ms following fixation onset. In Fig. 4e–f, time-resolved RFs were inferred for responses aligned to saccade onset in 50-ms rolling response windows from -375–375 ms in 25-ms steps.

For saccade-aligned RFs, the above process was repeated for two retinotopic coordinate frames, anchored on the fixation point either before (FP 1) or after (FP 2) the saccade. Saccades were selected to be at least 5 dva in size to reduce stimulus feature correlation. Still, there may be residual spurious correlations due to finite saccade sizes and autocorrelations in natural images. To quantify the empirical baseline RF, we calculated, as a control, a third set of RFs anchored on the midpoint passed by the saccade.

The process above specifies the quantification described in the main text. In the supplementary material, we show implementation detail variants, which do not change the main text conclusions.

#### Simulation of responses representing ground-truth RFs

Each simulated RF was discretized into one or more offset locations in 2-dva steps, to be indexed into corresponding 2 × 2 image patches aligned to each eye position sample. Offset locations were assigned weights based on a Gaussian decay profile truncated at σ = 2. Responses were simulated for each eye position sample at its native 1 kHz sampling rate, although downstream analysis would bin responses into 50-ms time bins. A simulated response sample was the weighted sum of the model representations of image patches. No stochasticity was added. The simulated responses were entered into the same analysis pipeline as described above for real data. To prevent trivial generalization from the neural network representations (ViT) underlying the RF inference analysis, the simulated responses were based on model embeddings in ResNet-50 (He et al., 2016) (implementation and ImageNet-pretrained weights from the Python library ‘torchvision’) at the layer ‘layer3.15.bn2’.

#### Average estimates and statistical tests

In Fig. 2c and e, normalized responses were averaged over neurons. The spread was reported as the standard deviation over neurons. In Figs. 1g, 3f–h, 3k, 4b, 4c, and 4g, the metrics were averaged first over neurons per monkey, then over monkeys. The spread was reported as the 95%-confidence interval of the mean by bootstrapping first over neurons, then over monkeys. The statistical tests in Figs. 2c, 2e, and 4g were one-tailed paired permutation tests. P-values were corrected for the false discovery rate at 0.01 using the two-stage Benjamini-Krieger-Yekutieli procedure.

## Extended Data Figures

**Extended Data Fig. 1.**
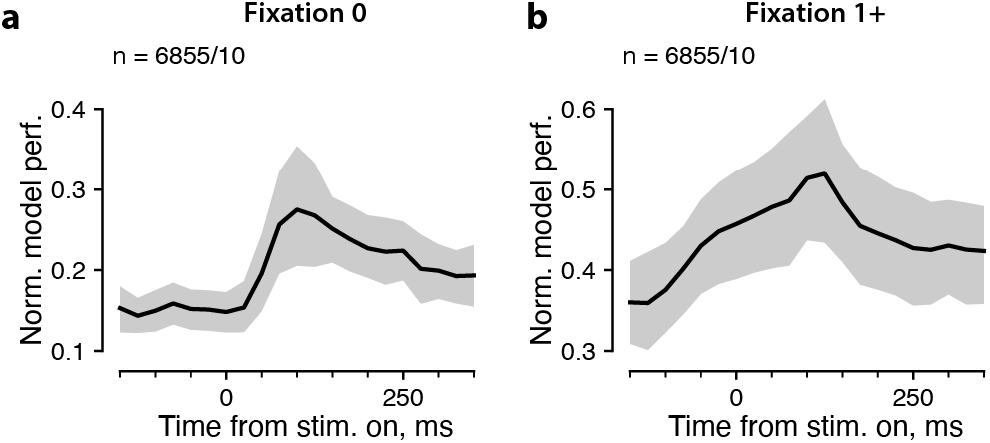
Computational models captured a large fraction of explainable individual-fixation responses. We normalized the cross-validated model performance by the ceiling of return fixation self-consistency per neuron. Panels **a** and **b** respectively show zeroth fixations and subsequent fixations. Plotting conventions follow that in Fig. 4b, c.

**Extended Data Fig. 2.**
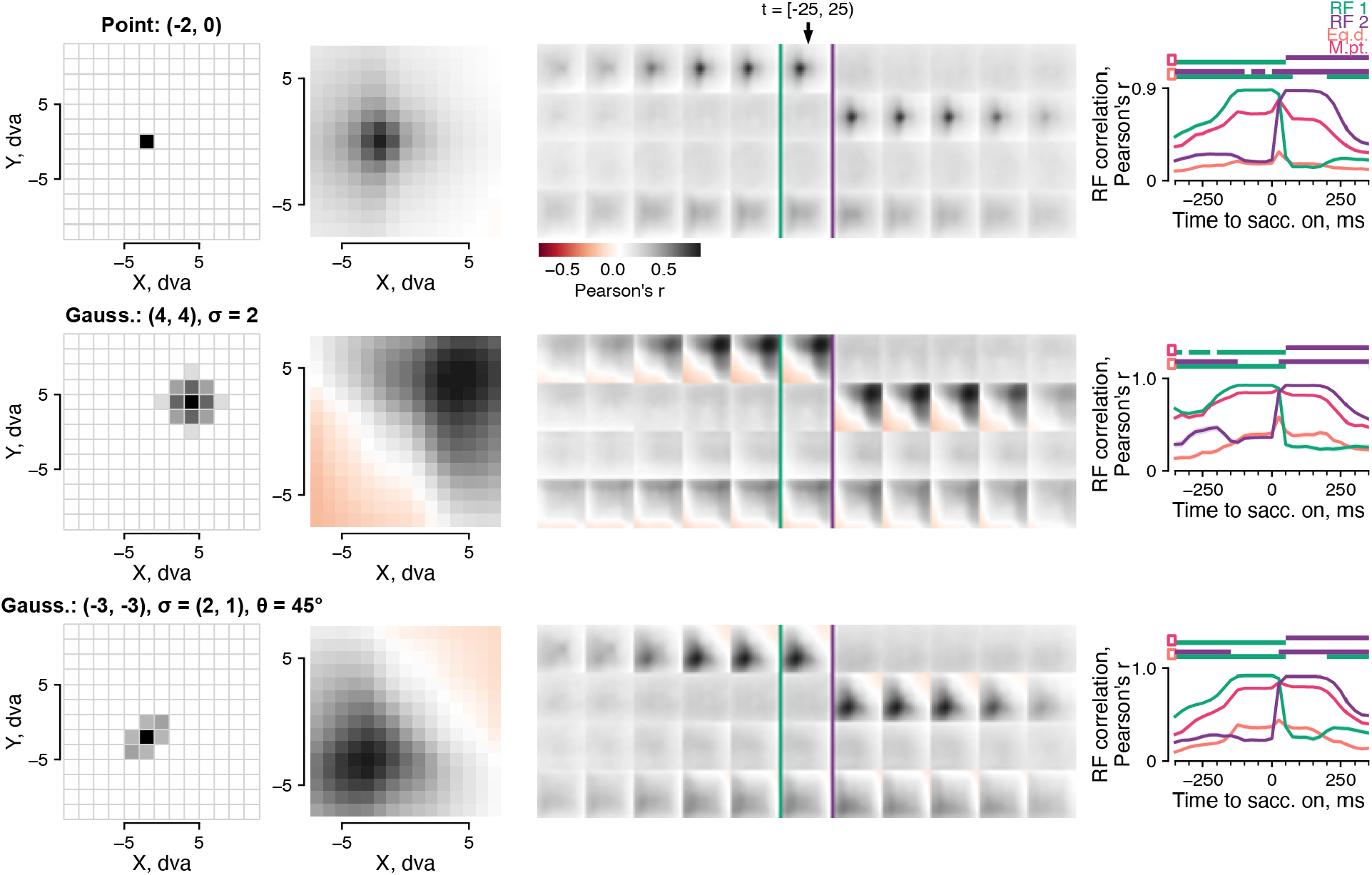
Validation of RF inference analysis using simulated activity. We simulated retinotopic, stimulus-selective activity using example behavior data from one session, then submitted the simulated responses to the RF inference analysis (Fig. 4d). The spatial profile of the responses was simulated using a weighted combination of ResNet-50 representations of 2 × 2 image patches on a retinocentric grid. The simulated responses had zero temporal lag relative to the eye position. (See Methods for details.) Each row corresponds to a different simulated RF. The first column illustrates the image patches and weights used to approximate the RF. The second column corresponds to Fig. 4e. In the third column, the first two rows of each set of maps correspond to Fig. 4f. The third and fourth rows correspond to the two counterfactual controls, equidistant (row 3) and midpoint (row 4). The arrow and purple/green lines indicate the response time bin -25–25 ms after saccade onsets. The fourth column corresponds to Fig. 4g.

**Extended Data Fig. 3.**
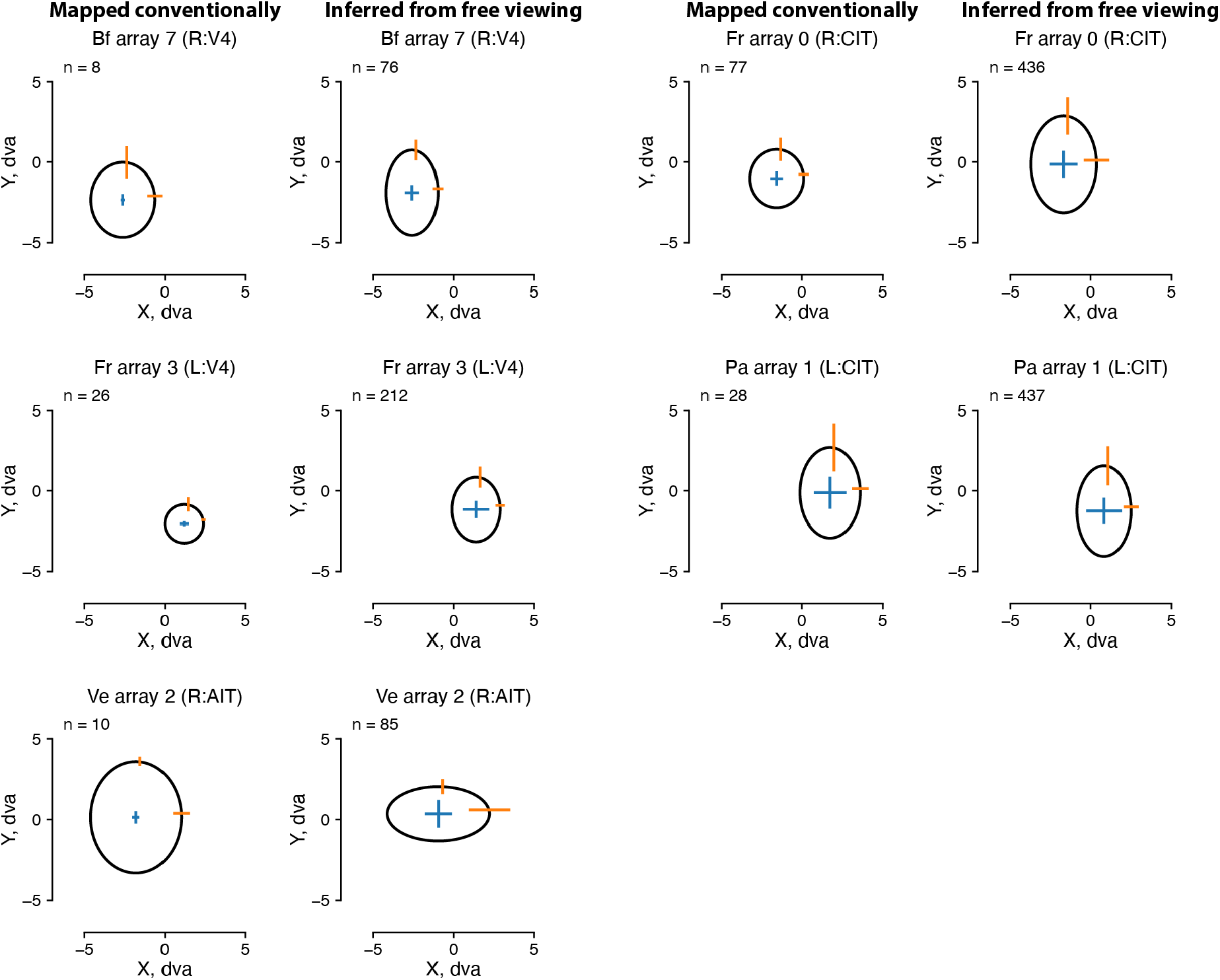
Model-inferred free viewing RFs matched conventionally mapped ones. RFs recovered from traditional mapping data (firing rates as a function of location in response to small stimuli flashed at various locations) were compared to model-inferred RFs (Fig. 4d–g). In each subplot, the ellipse and error bars indicate the median ± median absolute deviation of the center and size of RF Gaussian fits across neurons within an array. The five pairs of plots show five example arrays; titles indicate the monkey, hemisphere, and visual area of the arrays. Columns 1 and 3 correspond to conventionally mapped RFs; columns 2 and 4 correspond to model-inferred RFs from free viewing data.

**Extended Data Fig. 4.**
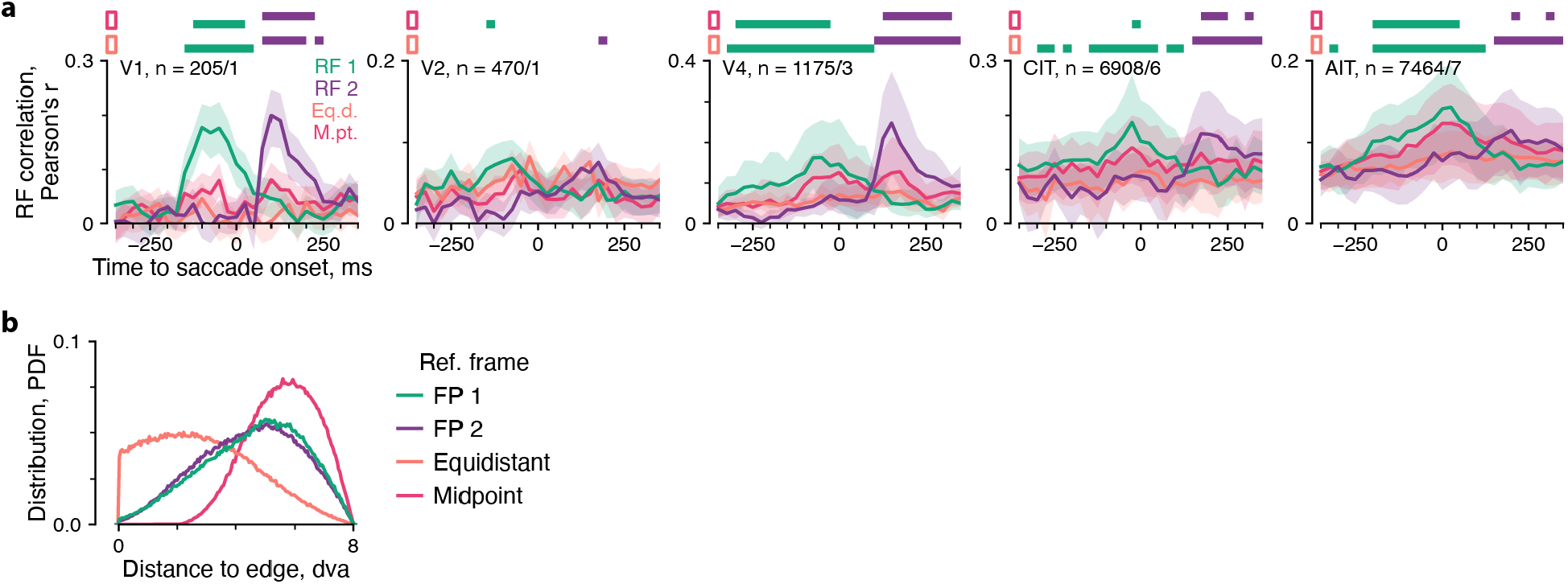
RFs quantified by fit to zeroth fixation and statistics of control reference frames for mapping. **a**, Quantification of RF presence over time and reference frames by consistency to zeroth-fixation RFs. Plotting conventions follow Fig. 4g. **b**, Distribution of center distance to image edge for the different mapping reference frames.

**Extended Data Fig. 5.**
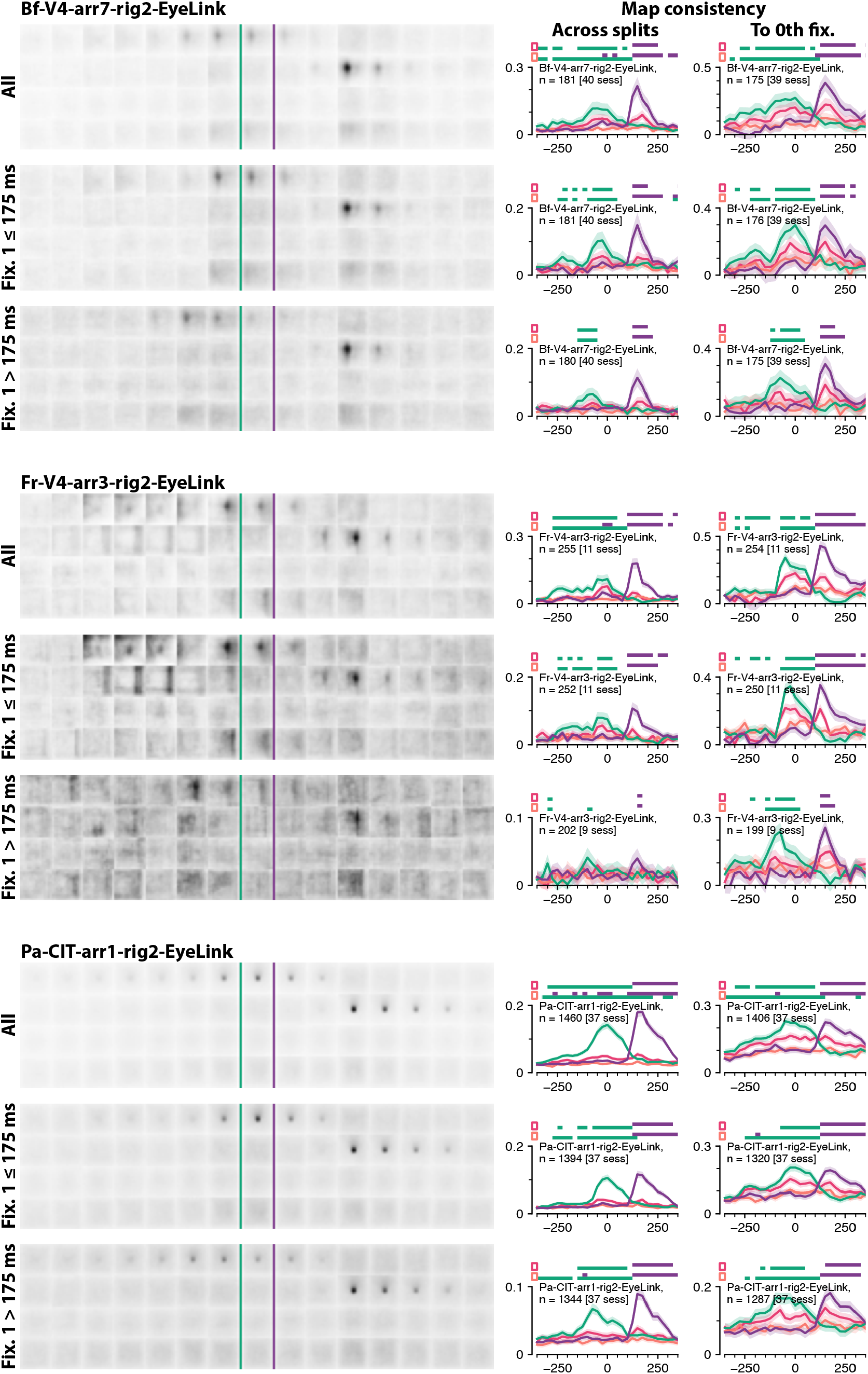
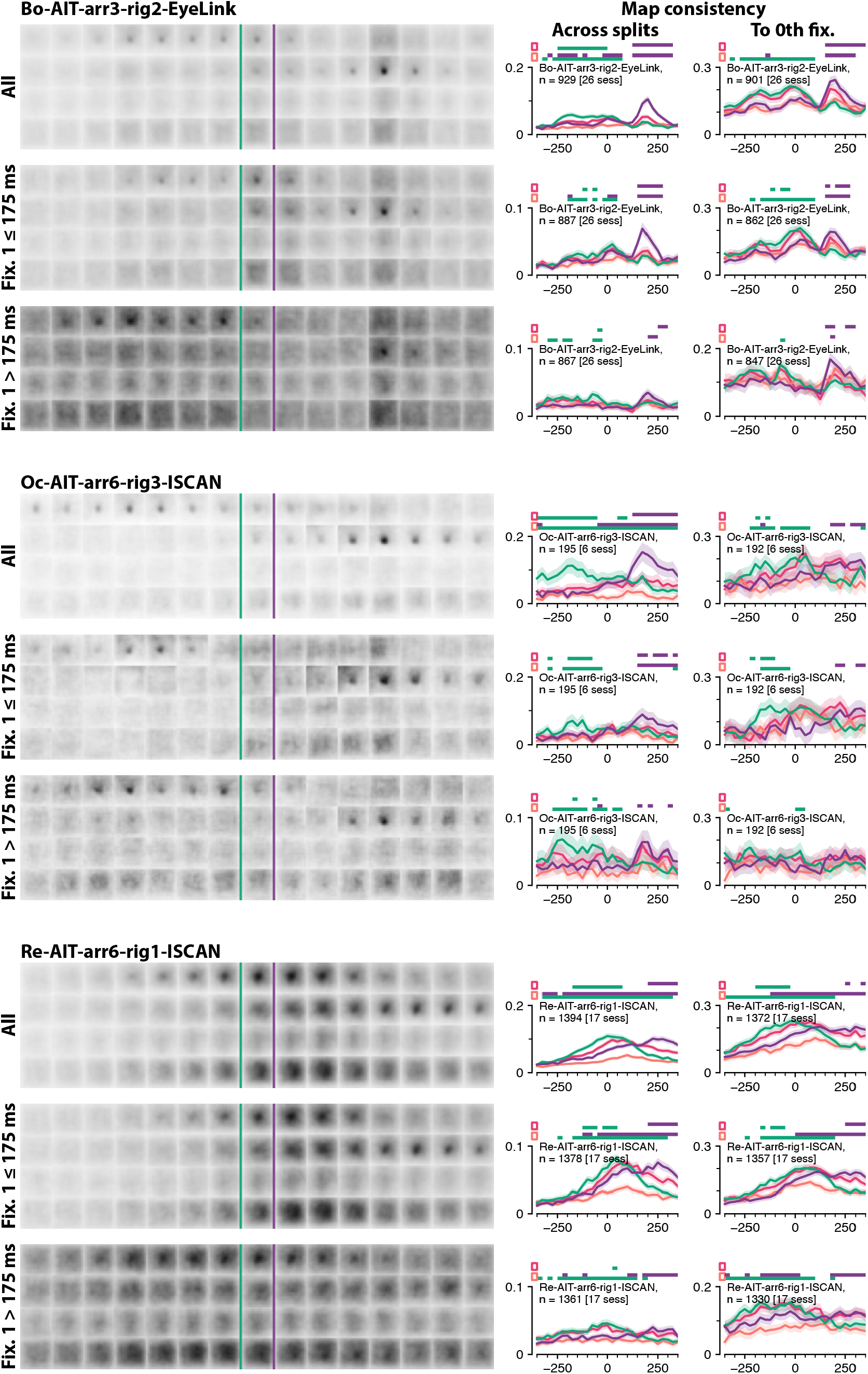
Example RFs that showed evidence for predictive remapping reaching statistical significance. Each group of three rows corresponds to data from an array collected with a specific experimental rig and eye tracker. Within each group, rows 2 and 3 correspond to mapping done on subsets of saccades that started with either a short- or a long-duration first fixation, to evaluate whether the predictive effect was aligned in time to the first fixation onset instead of saccade onset (see the main text for details). Within each row, the first column visualizes the inferred RFs averaged across neurons following Fig. 4f and Extended Data Fig. 2. Columns 2 and 3 quantify the RFs as in Fig. 4g.

